# The GBAF chromatin remodeling complex binds H3K27ac and mediates enhancer transcription

**DOI:** 10.1101/445148

**Authors:** Kirill Jefimov, Nicolas Alcaraz, Susan L. Kloet, Signe Värv, Siri Aastedatter Sakya, Christian Dalager Vaagenso, Michiel Vermeulen, Rein Aasland, and Robin Andersson

**Affiliations:** Department of Biological Sciences, University of Bergen, 5008 Bergen, Norway; The Bioinformatics Centre, Department of Biology, University of Copenhagen, 2200 Copenhagen, Denmark; Department of Molecular Biology, Radboud Institute for Molecular Life Sciences, Radboud University Nijmegen, 6525 Nijmegen, The Netherlands; Department of Biosciences, University of Oslo, 0371 Oslo, Norway

## Abstract

H3K27ac is associated with regulatory active enhancers, but its exact role in enhancer function remains elusive. Using mass spectrometry-based interaction proteomics, we identified the Super Elongation Complex (SEC) and GBAF, a non-canonical GLTSCR1L- and BRD9-containing SWI/SNF chromatin remodeling complex, to be major interactors of H3K27ac. We systematically characterized the composition of GBAF and the conserved GLTSCR1/1L ‘GiBAF’-domain, which we found to be responsible for GBAF complex formation and GLTSCR1L nuclear localization. Inhibition of the bromodomain of BRD9 revealed interaction between GLTSCR1L and H3K27ac to be BRD9-dependent and led to GLTSCR1L dislocation from its preferred binding sites at H3K27ac-associated enhancers. GLTSCR1L disassociation from chromatin resulted in genome-wide downregulation of enhancer transcription while leaving most mRNA expression levels unchanged, except for reduced mRNA levels from loci topologically linked to affected enhancers. Our results indicate that GBAF is an enhancer-associated chromatin remodeler important for transcriptional and regulatory activity of enhancers.

**Graphical abstract:** 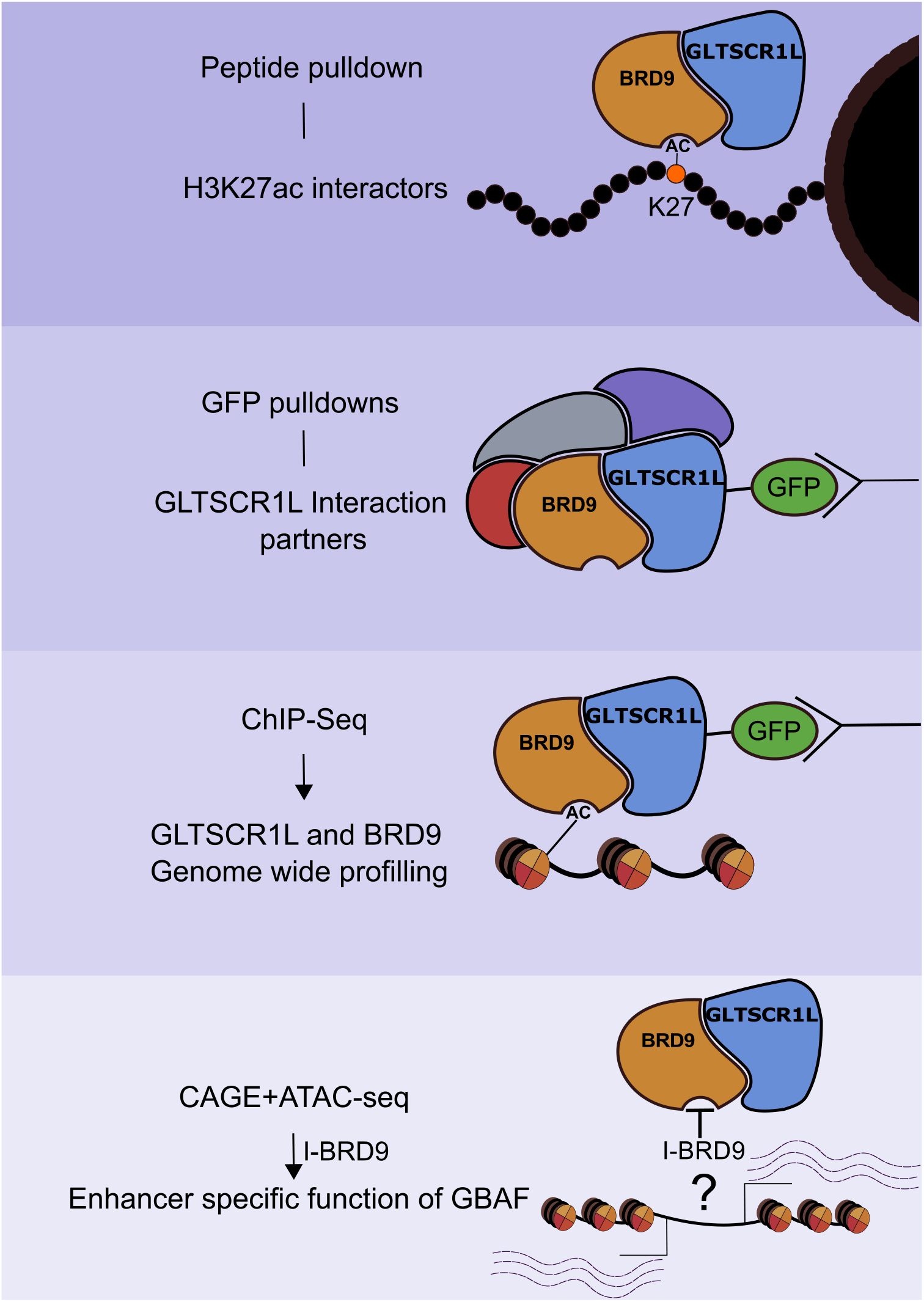

---

Regulatory events at gene promoters and transcriptional enhancers modulate cell-type specific gene activities and allow cells to respond to external cues (Heinz et al. 2015; Beagrie & Pombo 2016; Lenhard et al. 2012). Several processes take part in these actions, including ATP-dependent chromatin remodeling, transcription factor (TF) and co-activator binding, and the recruitment of general transcription factors (GTF) and RNA polymerase II (Pol II) (Vernimmen & Bickmore 2015). Integral to the activation of gene transcription is a favorable chromatin environment. Gene transcriptional activity is associated with permissive histone post-translational modifications (PTMs) (Li et al. 2007) and a three-dimensional folding of the genome that brings enhancers in close proximity with gene promoters (Sanyal et al. 2012; Rao et al. 2014), thereby allowing the regulation of target genes.

Genome-wide charting of histone PTMs has identified chromatin signatures associated with repressive and transcriptionally permissive states as well as enhancer activity (Jenuwein 2001; Heintzman et al. 2009; Kundaje et al. 2015). However, the mechanisms by which many histone PTMs exert their putative functions are elusive. These may include the modulation of nucleosome-DNA interaction strength or recruitment of protein complexes that execute specific functions (Kouzarides, 2007). Among many potential histone PTMs, acetylation of histone H3 at lysine 27 (H3K27ac) is associated with regulatory activity of both gene promoters and enhancers (Creyghton et al., 2010, Ernst et al., 2011, Rajagopal et al., 2014, Rada-Iglesias et al., 2011). H3K27ac is deposited by acetyltransferases CBP and P300 (Tie et al. 2009). Targeting of P300 fused with nuclease-null dCas9 has been reported to elevate H3K27ac levels and activate transcription from promoter-proximal as well as distal regulatory regions (Hilton et al. 2015).

Although the association of H3K27ac with active regulatory elements is known, its potential role in enhancer function is still unclear. Given its recognition by acetyl-lysine reader domains, such as BROMO and YEATS, it is possible that H3K27ac attracts effector proteins to enhancers. Previous work to identify H3K27ac-interacting proteins has relied on chromatin immunoprecipitation followed by identification of co-purified proteins by mass-spectrometry (ChIP-MS) (Engelen et al. 2015; Ji et al. 2015). Unfortunately, due to experimental setup (purification of sheared, crosslinked chromatin with pre-existing protein complexes), it is hard to discern the contribution of H3K27ac to enhancer function from DNA-dependent activities, such as TF binding.

Here, we applied a SILAC-based histone-peptide pulldown approach to identify proteins interacting with H3K27ac in mouse embryonic stem cells and HeLa cells, also in combination with H3K23ac. These experiments revealed Super Elongation Complex (SEC) and BRM/BRG1 Associated Factors (BAF) as major interactors of H3K27ac. Furthermore, we identified GLTSCR1L protein to be a part of a H3K27ac-specific BAF complex, recently identified as GBAF (Alpsoy & Dykhuizen 2018), and we characterized the interaction partners of the GLTSCR1L protein and its conserved protein-interaction domain. Our findings led us to further investigate the function of the GLTSCR1L-BRD9 containing GBAF complex in transcription. Upon treatment with a BRD9 bromodomain inhibitor, we observed enhancer-specific transcriptional abnormalities, indicating an important role for GBAF in enhancer transcription and regulatory activity.

## Results

### SWI/SNF and Super Elongation Complex are major acetyl-lysine interacting complexes

We applied an established SILAC histone peptide pulldown approach (Vermeulen et al. 2007) to identify proteins that specifically bind to H3K27ac (Methods). To discriminate proteins that bind specifically to H3K27ac from proteins with a general affinity for acetyl-lysine-containing histone peptides, we performed the same experiment with peptides associated with the highly abundant (Sidoli et al. 2015; Tvardovskiy et al. 2015) H3K23ac mark (Fig. 1a,b). In addition, we included an experiment with histone peptides diacetylated at both H3K23 and H3K27 residues (Fig. 1e,f), since these acetylations frequently colocalize and are associated with active transcription (Wang et al. 2008). To understand how the recognition of H3K27ac compares across mammals and whether there are any differences between pluripotent and terminally differentiated states, we used both HeLa cells and mouse embryonic stem cells (mESCs) as sources of nuclear extracts for the pulldown experiments. Each peptide pulldown experiment resulted in the identification of more than 2,000 proteins, of which 10 to 30 showed significant acetyl-lysine specific binding (Fig. 1a-c, Supplementary Fig. 1a-c, Supplementary Data 1). Our experimental design allowed us not only to identify acetyl-lysine binding complexes, but also to estimate their binding preferences (H3K27ac versus H3K23ac). We averaged forward and reverse SILAC ratios of each individual pulldown separately and applied hierarchical clustering. Proteins from the same complexes were found to cluster together (Fig. 1d, Supplementary Fig. 1d). The majority of identified histone H3 acetyl-lysine interacting proteins are members of two distinct protein complexes: the SEC and the SWI/SNF family of chromatin remodeling complexes (Wang et al. 1996; Lin et al. 2010). In addition, we detected NuA4 complex components DMAP1 and YEATS4 (Doyon et al. 2004) and components of the general transcription factor TFIID (Dynlacht et al. 1991).

**Figure 1.**
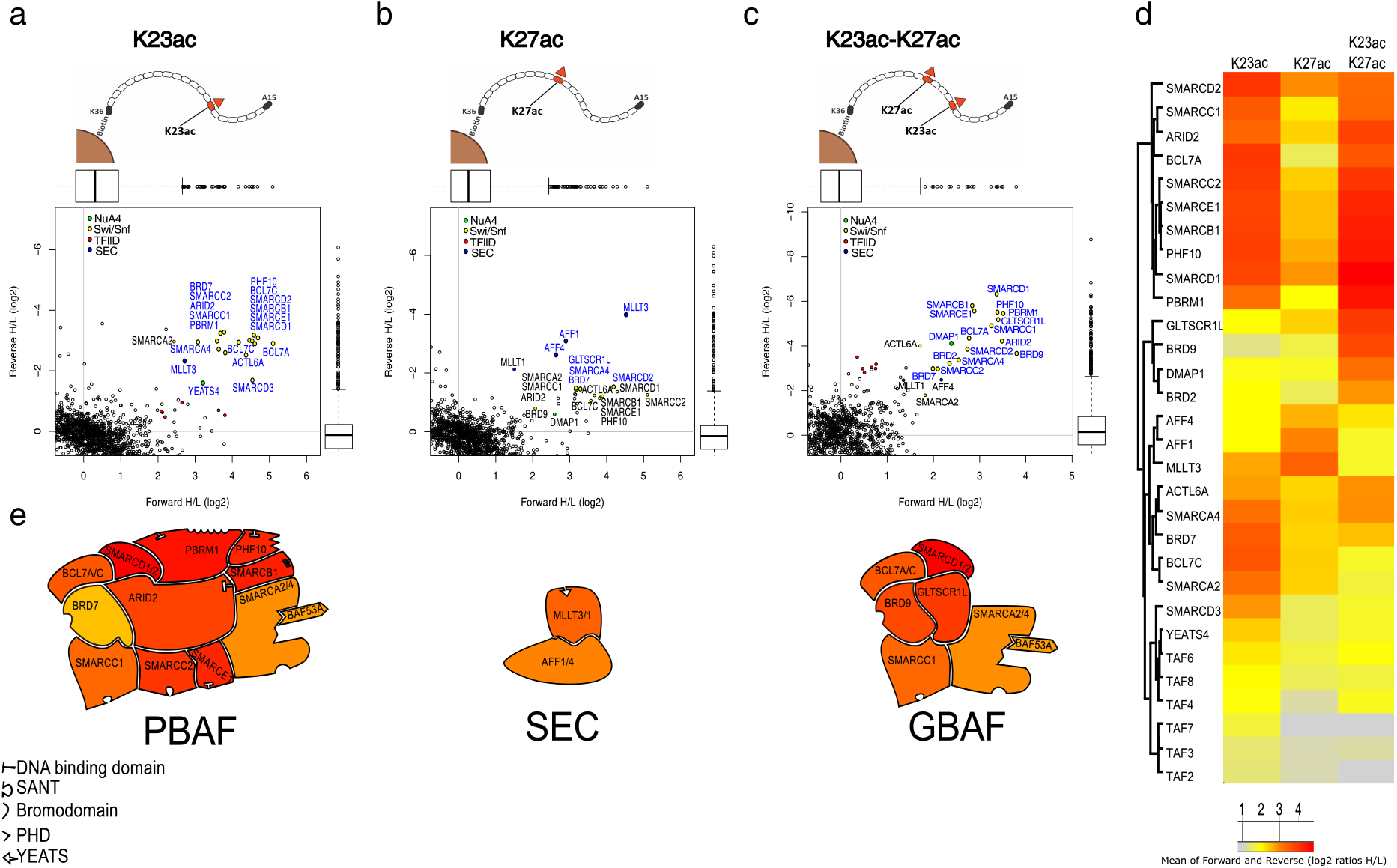
SILAC histone peptide pulldowns identify SEC and SWI/SNF complexes as major interactors of H3K23ac and H3K27ac. **a-c** Scatter plots of forward (horizontal axes) versus reverse (vertical axes) SILAC ratios from histone peptide pulldowns using (**a**) H3K23ac, (**b**) H3K27ac, or (**c**) diacetylated H3K23acK27ac baits with HeLa cell nuclear extracts (n=2 pulldowns for each bait). The lower and upper hinges of boxes correspond to the first and third quartiles of data, respectively, and the whiskers extend to the largest and smallest data points no further away than 1.5 times the interquartile range. **d** Heatmap of average forward and reverse SILAC ratios (log_2_). The ordering of rows in the heatmap and the associated dendrogram were derived from agglomerative hierarchical clustering of the H3K23- and K27-interacting proteins identified in the histone peptide pulldowns (**a-c**), using Euclidean distances. **e** Visual summary of the most enriched interactors of H3K23ac (left), H3K27ac (middle), and H3K23acK27ac (right) peptide pulldowns. Subunits are colored according to the log2 value of their enrichment in the pulldowns.

The SEC complex is important for the activation of transcription by release of paused Pol II (Lin et al 2010; Luo et al, 2012). Although many SEC complex components were identified in pulldowns with all acetylated histone peptides, we observed differences in SILAC ratios between nuclear extracts. The highest SILAC ratio for the SEC complex subunits was observed with monoacetylated H3K27ac histone peptide with HeLa and diacetylated H3K23acK27ac peptide with mESC nuclear extracts. In HeLa cells, we were able to detect the presence of AF9, AFF4, AFF1 and ENL, but not ELL family proteins nor P-TEFb (Fig. 1b), important mediators of SEC and Pol II interaction (Luo et al, 2012; Knutson et al, 2016). In mESCs, we detected nearly all SEC complex subunits, including AF9, AFF4, ELL3 and CDK9 (Supplementary Fig. 1c). AF9 contains a YEATS-domain that is able to recognize histone modifications and has previously been found to bind H3K9ac and H3K27ac (Li et al, 2014).

Mammalian SWI/SNF (or BAF) complexes are ATP-dependent chromatin remodelers implicated in a wide variety of processes in the cell nucleus (Hargreaves & Crabtree 2011). Different combinations of SWI/SNF subunits generate a diversity of functionally distinct complexes, including two canonical subfamilies (BAF, PBAF), their subcomplexes, and GBAF, a recently described non-canonical SWI/SNF complex (Phelan et al. 1999; Hohmann & Vakoc 2014; Alpsoy & Dykhuizen 2018). We observed known core SWI/SNF subunits (SMARCA2/4, SMARCC1, SMARCC2) and shared subunits (SMARCB1, SMARCE1, SMARCD1/2/3, ACTL6A, BCL7A/C) in pulldowns with each of the acetylated histone peptide baits, with both HeLa and mESC nuclear extracts. The complete PBAF complex, represented by ARID2 (BAF200), PHF10 (BAF45A), PBRM1 (BAF180) and BRD7, was found to prefer interaction with H3K23ac-containing peptides (both mono-acetylated H3K23ac and di-acetylated H3K23acK27ac peptides) (Fig. 1a,c, Supplementary Fig. 1a,c). However, known BAF complex specific subunits (ARID1A/1B (BAF250A/B), DPF1/2/3 (BAF45B/C/D)) were absent from these pulldowns. Instead, we observed BRD9 and GLTSCR1L (BICRAL)^1^ proteins, subunits of the GBAF complex (Alpsoy & Dykhuizen 2018). These proteins predominantly interact with H3K27ac-containing peptides. GLTSCR1L was the most enriched interactor of monoacetylated H3K27ac in mESCs and one of the most enriched H3K23acK27ac interactors in HeLa cells (Fig.1c, Supplementary Fig. 1b).

Taken together, mass spectrometry-based interaction proteomics experiments identified SWI/SNF and SEC as major acetyl-lysine readers, which is consistent with earlier findings (Li et al. 2014; Chandrasekaran & Thompson 2007; Chandy et al. 2006). Additionally, we identified GBAF components GLTSCR1L and BRD9 as prominent H3K27ac-interacting proteins.

### GLTSCR1L is a subunit of a distinct non-canonical BAF complex

Due to the observed preferential binding of the GLTSCR1L protein to H3K27ac-containing histone peptides, we decided to identify GLTSCR1L-interacting proteins by quantitative interaction proteomic methods. We generated a HeLa cell line stably expressing doxycycline-inducible GFP-tagged GLTSCR1L and performed label-free GFP pulldowns followed by liquid chromatography-MS (LC-MS/MS, Methods) (Smits et al. 2013). The full length GFP-GLTSCR1L protein interacted with core SWI/SNF subunits including SMARCD1 (BAF60a), SMARCC1 (BAF155), SMARCA4 (BRG1), and BAF-specific subunit SS18 (Fig. 2a). One of the strongest GFP-GLTSCR1L interactors was BRD9, which also clustered together with GLTSCR1L in our histone peptide pulldown analysis (Fig. 1d, Supplementary Fig. 1d). Additionally, BCL7C was identified as a novel interactor of GLTSCR1L, while other canonical BAF complex subunits, such as ARID1A, DPF1/2/3 (BAF45B/C/D), and SMARCE1 (BAF57), were absent in the pulldowns.

**Figure 2.**
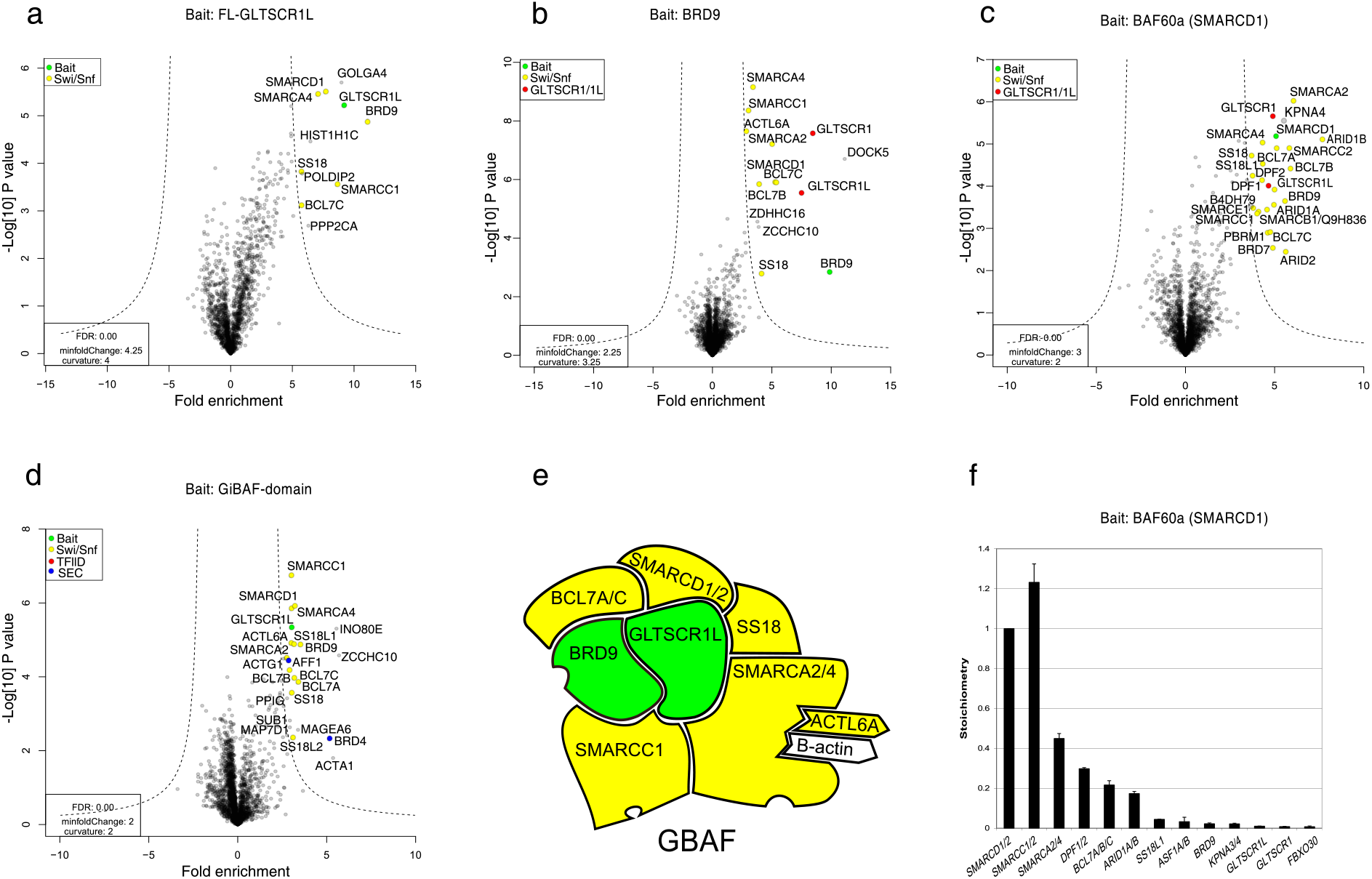
Quantitative MS analysis reveals the interacting proteins of GLTSCR1L protein and its GiBAF-domain. **a-d** Volcano plots displaying fold enrichments (log_2_ GFP fusion protein over mock, horizontal axes) versus t-test P values (-log_10_, vertical axes) of identified interactors of affinity-purified GFP-GLTSCR1L (**a**), GFP-BRD9 (**b**), GFP-BAF60a (SMARCD1) (**c**) and GFP-GiBAF-domain of GLTSCR1L (**d**) isolated from HeLa whole cell extracts (**a-c**) or nuclear extracts (**d**). Each volcano plot represents three independent pulldowns. **e** Schematic representation of the GBAF complex. Proteins in green have been used as baits, proteins in yellow represent identified interactors. **f** Stoichiometry of GFP-BAF60a interactors. The intensity-based absolute quantification (iBAQ) value of each protein is divided by the iBAQ value of the bait and plotted relative to a bait value of 1. Data are shown as mean ± s.d. (*n* = 3 pulldowns).

In order to validate GLTSCR1L as a *bona fide* interactor of SWI/SNF complexes, we performed reciprocal label-free GFP pulldowns. We generated three doxycycline-inducible HeLa cell lines expressing GFP-tagged baits of identified GLTSCR1L-interacting proteins: BRD9 (Fig. 2b), SMARCD1 (BAF60a) (Fig. 2c) and SMARCC1 (BAF155) (Supplementary Fig. 3a). As expected, we observed an enrichment of SWI/SNF complex subunits in all pulldowns. The highest number of identified SWI/SNF subunits was observed in pulldowns with SMARCC1 (BAF155) as a bait, followed by SMARCD1 (BAF60a) and BRD9 pulldowns (Supplementary Table 1). Additionally, GLTSCR1L was detected as an interactor of each of these three bait proteins (Fig. 2b,c, Supplementary Fig. 3a). We also observed a nearly complete overlap of interactors in the GFP-BRD9 and GFP-GLTSCR1L pulldowns (Fig. 2a,b), suggesting that these two proteins frequently co-occur in the same sub-complex. Interestingly, the homologous protein GLTSCR1 also appeared in all reciprocal pulldown experiments but not in pulldowns with GLTSCR1L, indicating that GLTSCR1 and GLTSCR1L proteins bind to SWI/SNF complexes in a mutually exclusive manner.

The observed binding of both GLTSCR1 and GLTSCR1L to SWI/SNF complexes led us to investigate the role of their shared, conserved domain in mediating the interaction with SWI/SNF family members. We refer to the domain as ‘GiBAF’, for <u>G</u>LTSCR1/1L domain interacting with <u>BAF</u> complex. First, we produced a TY1-tagged GLTSCR1L protein containing an in-frame deletion of the GiBAF-domain. By means of immunofluorescence microscopy (IF) with FL and domain deletion mutant proteins, we found that the GiBAF-domain is responsible for the nuclear localization of GLTSCR1L (Supplementary Fig. 3b). Next, we generated two other HeLa cell lines with doxycycline-inducible GFP-tagged GLTSCR1L protein mutants expressing only the GiBAF-domain or the full-length protein lacking the domain (▲GiBAF-domain). In pulldowns with the GiBAF-domain only, we observed all full length GFP-GLTSCR1L protein interactors in addition to some other BAF complex components (BCL7A/B/C, ACTL6A, SMARCA2, SS18/L1/L2) (Fig. 2d). GiBAF-domain interaction with BAF155, BAF60a and BRD9 was also confirmed by immunoblotting (Supplementary Fig. 3g) (Alpsoy & Dykhuizen 2018). Additionally, we detected an interaction between the GiBAF-domain from GLTSCR1L protein and BRD4, in line with previous work demonstrating an interaction between GLTSCR1 and the BRD4 ET-domain, which contributes to transcriptional regulation of BRD4 target genes (Rahman et al. 2011). We also observed interaction with INO80E (Chen et al. 2011) from the INO80 (Jin et al. 2005) chromatin remodeling complex, which has previously been shown to interact with SWI/SNF complex components (Cai et al. 2007; Yao et al. 2008). AFF1, a SEC complex subunit, was also found to interact with the GiBAF-domain of GLTSCR1L. Similar analysis performed with the GiBAF-domain deletion mutant of GLTSCR1L protein (▲GiBAF-domain) resulted in no significant interactions with BAF complex subunits (Supplementary Fig. 3c), further supporting a role for the GiBAF-domain in mediating interactions with other SWI/SNF subunits. Instead, we observed a significant enrichment of TAF6 and TAF4 TFIID components co-purified with ▲GiBAF-domain, indicating that other parts of the GLTSCR1L protein may be responsible for non-SWI/SNF protein interactions.

Label-free GFP pulldowns allowed to use the intensity-based absolute quantification (iBAQ) algorithm (Schwanhäusser et al. 2011) to calculate the relative abundance (stoichiometry) of observed proteins. Stoichiometry values from the FL GLTSCR1L and GiBAF-domain pulldowns (Supplementary Fig. 3d-e) confirmed a strong association between GLTSCR1L and BRD9. Interestingly, analyses of reciprocal pulldown-MS with BAF complex components SMARCD1 and SMARCC1 revealed very low levels of GLTSCR1L protein abundance (Fig. 2f, Supplementary Fig. 3f; ~1,6% of SMARCD1 and 0,066% of SMARCC1 are found together with GLTSCR1/1L), indicating that GLTSCR1L is either a transient interactor or a component of a rare BAF complex in HeLa cells. Taken together, these results suggest that GLTSCR1L is a subunit of a distinct SWI/SNF chromatin remodeling subcomplex, whose interactions with SWI/SNF subunits and its nuclear localization is mediated by the GiBAF-domain.

### Selective inhibition of the BRD9 bromodomain leads to disassembly of GLTSCR1L but not BRD9 from chromatin

Since the GLTSCR1L protein does not contain any known DNA/chromatin-binding domain, we hypothesized that, due to the interaction between GLTSCR1L and BRD9, the latter might facilitate targeting of GBAF to chromatin. In order to validate the association between GLTSCR1L, BRD9 and H3K27ac, we performed histone-peptide pulldown assays using I-BRD9 (Theodoulou et al. 2016), a specific inhibitor of the BRD9 bromodomain. I-BRD9 does not impair any other function of BRD9 beyond its acetyl-lysine recognition, making it a suitable tool to investigate the role of H3K27ac binding by the BRD9 bromodomain with limited secondary effects. Both GLTSCR1L and BRD9 were found to follow the trend of acetyl-lysine recognition observed in histone-peptide pulldown MS (H3K23ac<H3K27ac<H3K23acK27ac) (Fig. 1d). After treatment with I-BRD9, we detected a loss of specificity of BRD9 binding to H3K27ac-containing peptides. Although BRD9 still showed interaction with histone peptides while its bromodomain was blocked by I-BRD9, GLTSCR1L binding to H3K27ac-containing histone peptides was completely abrogated (Fig 3a).

**Figure 3.**
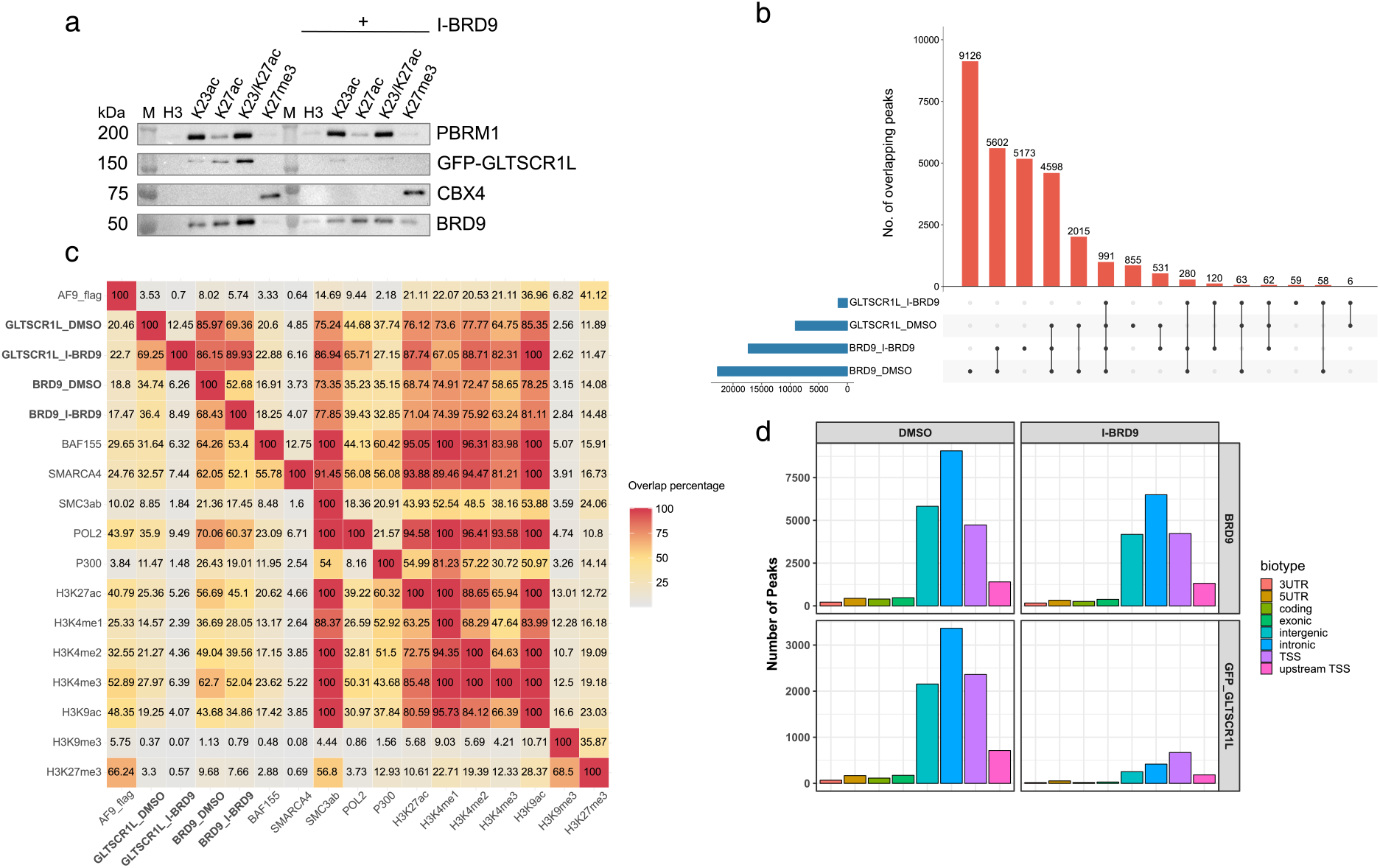
Inhibition of the bromodomain of BRD9 leads to disassembly of GLTSCR1L from chromatin. **a** Histone peptide pulldown assay using biotinylated histone H3 peptides (aa 15-36) and nuclear extract from HeLa-FRT-GFP-GLTSCR1L cells in the absence and presence of BRD9 bromodomain inhibitor I-BRD9. Unmodified (H3) and modified peptides were used. Input and affinity purified fractions were analyzed after SDS-PAGE by immunoblotting. PBRM1 and CBX4 serve as controls for I-BRD9 specificity. M: protein molecular weight standard. **b** Histogram attribute (upset) plot of ChIP-seq peak intersections between GLTSCR1L and BRD9 with and without I-BRD9. Vertical axis gives the frequency of overlaps between combinations of peak calls specified on the horizontal axis. **c** Heatmap of the overlap percentages between GLTSCR1L and BRD9 ChIP-seq peaks (bold) and ENCODE-called peaks of factors and chromatin marks associated with transcription. Each cell shows the percentage of ChIP-seq peaks for factors and marks in rows overlapping with ChIP-seq peaks for factors and marks in columns. **d** Genomic annotation of GLTSCR1L and BRD9 ChIP-seq peaks based on GENCODE-inferred (v19) genomic annotation biotypes.

To validate our histone-peptide pulldown results and to investigate whether GLTSCR1L, BRD9 and H3K27ac colocalize on chromatin, we performed chromatin IP of GLTSCR1L and BRD9 followed by high-throughput sequencing (ChIP-seq). We used a HeLa FRT cell-line expressing GFP-tagged GLTSCR1L and antibodies to GFP and wtBRD9, due to the unavailability of a ChIP-seq grade GLTSCR1L antibody. In parallel, we conducted similar ChIP-seq experiments with I-BRD9 treated cells. A total of 22,559 and 9,155 peaks were called (Irreproducible Discovery Rate < 0.05) for BRD9 and GFP-GLTSCR1L, respectively. We observed a strong colocalization between BRD9, GLTSCR1L, and H3K27ac (Supplementary Fig. 4a; permutation test, *P* < 1×10^−4^ for BRD9 and GLTSCR1L). In agreement with the histone-peptide binding assay, selective inhibition of the BRD9 bromodomain with I-BRD9 led to only a mild reduction of BRD9 binding to chromatin (22,559 versus 17,366 peaks for DMSO and I-BRD9, respectively), but severely abrogated GLTSCR1L chromatin binding (9,115 vs 1,639 peaks for DMSO and I-BRD9, respectively) (Fig 3b).

We observed a large degree of overlap of BRD9 and GLTSCR1L binding sites with ChIP-seq peaks of cohesin (SMC3ab) and active chromatin marks, especially H3K4me1/2, H3K27ac, and H3K9ac (Fig. 3c, 70-85% of GLTSCR1L and BRD9 peaks). GLTSCR1L and BRD9 was also associated with Pol II, with 70% of Pol II peaks overlapping with those of BRD9 and 35% with GLTSCR1L, suggesting a role for GBAF in the control of Pol II activity. Reassuringly, we detected chromatin colocalizations of both GLTSCR1L and BRD9 with core SWI/SNF subunits (~32% of BAF155 and SMARCA4 peaks overlapped with GLTSCR1L peaks and ~65% with BRD9). A small degree of colocalization was also observed between AF9 protein (a H3K9/K27ac reader), GLTSCR1L, and BRD9 (~20% of peaks for both proteins). However, only ~14% of GLTSCR1L and BRD9 peaks were found to overlap with H3K27me3 peaks compared to 41% for AF9, further pointing to a role of GBAF in transcriptionally active chromatin. Specifically, GLTSCR1L and BRD9 peaks were primarily located in intronic or intergenic regions and, to a lesser extent, at gene promoters. Upon I-BRD9 treatment, BRD9 largely remained at intronic/intergenic regions, whereas the greatly reduced binding sites of GLTSCR1L were mostly found at active gene TSSs (Fig. 3c,d). Taken together, our results indicate that GBAF binds active chromatin and that the bromodomain of BRD9 is responsible for GLTSCR1L binding to putative enhancers.

### BRD9 inhibition globally reduces enhancer transcription

To assess the role of GLTSCR1L and the GBAF complex in transcription, we examined I-BRD9 treated HeLa cells stimulated with epidermal growth factor (EGF) to induce a rapid transcriptional response. First, we inspected GLTSCR1L and BRD9 binding as well as RNA expression levels of the EGF-inducible gene *NR4A1* and its known enhancer 80 kb downstream of the *NR4A1* gene locus (Lai et al. 2015) by qPCR. EGF treatment increased binding of both GFP-GLTSCR1L and BRD9 proteins at the enhancer, but no increase was observed in I-BRD9 treated cells (Supplementary Fig. 5a). I-BRD9 did not affect the induction of *NR4A1* gene expression by EGF, in line with a non-significant enrichment of GFP-GLTSCR1L and BRD9 proteins at the *NR4A1* gene promoter, but almost completely abrogated expression of the enhancer RNA (Supplementary Fig. 5b).

We next performed 5’ end sequencing of capped RNAs, using Cap Analysis of Gene Expression (CAGE (Takahashi et al. 2012)), to assess the effect of EGF-induced transcription initiation events and enhancer activities (Andersson et al. 2014) genome-wide. CAGE experiments were performed in parallel with ATAC-seq across the EGF-response time course to focus on transcription initiation events at open chromatin loci such as enhancers and gene promoters. Using CAGE data in combination with ATAC-seq data, we were able to dissect the NR4A1 super-enhancer into a set of five bidirectionally transcribed enhancers in open chromatin with similar transcriptional EGF-response patterns, and found all of them to be downregulated by I-BRD9 (Fig. 4a, focusing on three enhancer constituents). Genome-wide, we next focused on all transcribed nucleosome-free regions (NFRs), inferred from ATAC-seq peaks associated with CAGE expression above estimated background noise level (Methods). Transcribed NFRs corresponded to ~11% (23,639 out of 213,017) of all detected open chromatin loci and displayed clear differences in expression patterns between I-BRD9 and DMSO-treated samples (52% of variance explained) as well as between time points (Supplementary Fig. 5c). Time points 30 and 60 minutes displayed different transcriptional activities than time points 0 and 240 minutes after treatment, indicating that at 240 minutes after EGF induction most transcriptional activities had returned to baseline levels.

**Figure 4.**
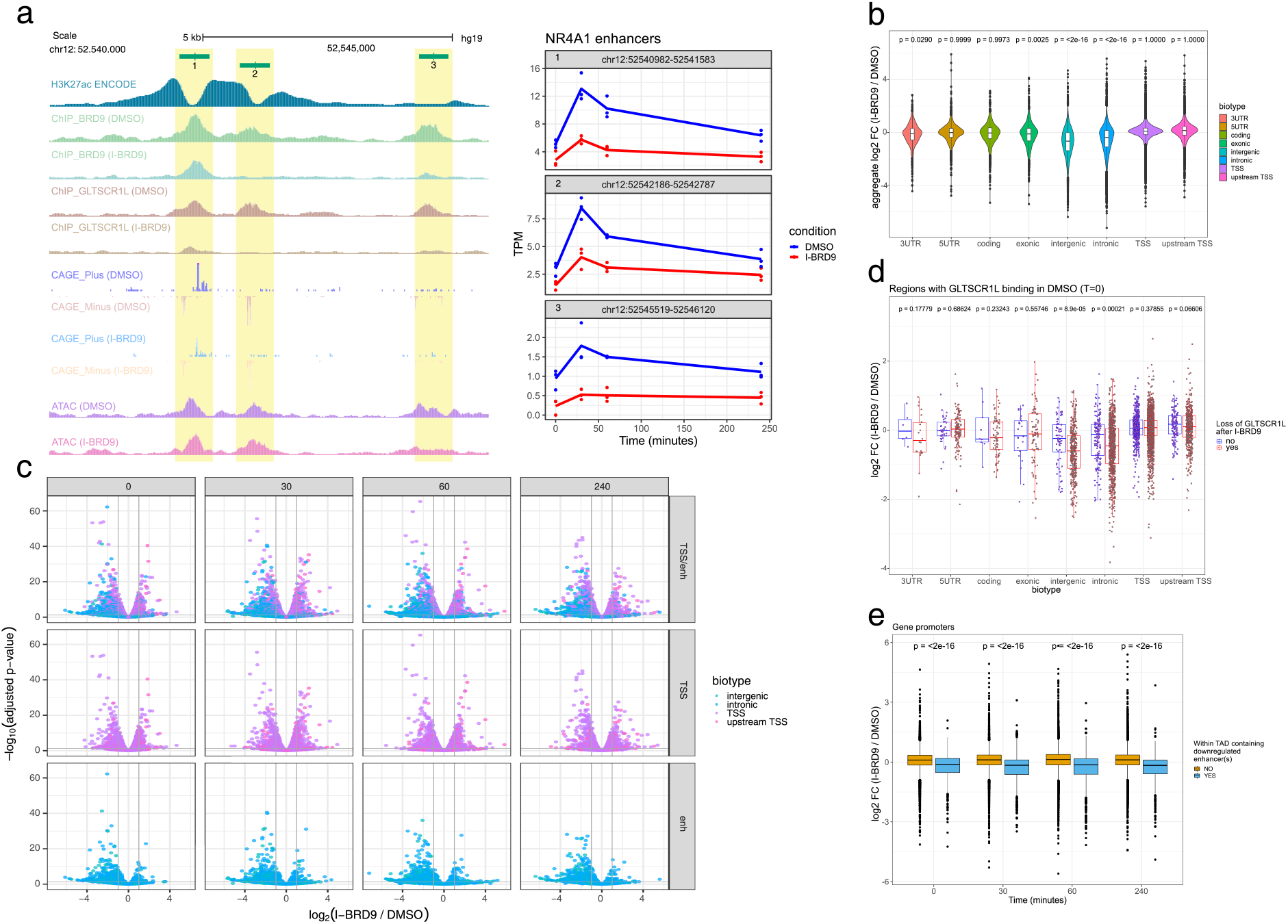
Inhibition of the bromodomain of BRD9 globally inhibits enhancer transcription. **a** Genome browser view (left) of the NR4A1 super-enhancer locus, showing tracks with pile-up signal from H3K27ac (ENCODE, HeLa S3), GLTSCR1L and BRD9 ChIP-seq, ATAC-seq, and CAGE data. Differences in ChIP-seq signal (except for H3K27ac), ATAC-seq signal, and CAGE expression between control (DMSO) and I-BRD9 treated cells are observable at three highlighted super-enhancer constituents before EGF stimulus. CAGE expression levels (TPM normalized) are shown (right) for the three enhancers across the HeLa EGF stimulation time course, comparing control (DMSO) and I-BRD9 treated cells. **b** Violin plots depicting densities of aggregated fold changes of expressed NFRs across the EGF treatment time course, broken up by genomic annotation biotype inferred from GENCODE (v19). **c** Volcano plots of NFR expression fold changes (horizontal axes) and adjusted P-values of differential expression (vertical axes, Methods), comparing I-BRD9 treated cells with non-treated (DMSO), broken by biotype and time point after EGF stimulation. **d** Box-and-whisker plot (as in Fig. 1) of expression fold changes (log_2_) comparing control (DMSO) and I-BRD9 inhibited cells at GLTSCR1L-bound expressed NFRs (before EGF stimulus) that lose or keep GLTSCR1L binding upon I-BRD9 treatment. **e** Box-and-whisker plot (as in Fig. 1) of fold changes (I-BRD9 versus control) of gene promoters within TADs that contain or do not contain down-regulated enhancers.

To investigate the effect of I-BRD9 on transcriptional responses, we compared the aggregated fold change of NFR expression levels in I-BRD9 treated cells and control cells across all EGF time points. Similar to our observation at the NR4A1 enhancer locus, the EGF responses of putative enhancers, as indicated by transcribed intergenic and intronic loci (75% and 25% of expressed intergenic and intronic regions overlap with FANTOM5 enhancers (Andersson et al. 2014), respectively), were significantly downregulated upon BRD9 inhibition (Wilcoxon signed-rank Test, lower tail, P < 2.2×10^−16^) in contrast to the responses of mRNA promoters, which were largely not affected (Fig. 4b). We further compared transcriptional events measured at each time point in control and I-BRD9 treated cells (Fig. 4c). Most downregulated events (log_2_ FC < -1, FDR adjusted *P* < 0.05) were detected at one hour after EGF induction (Supplementary Fig. 5d) and occurred mostly at intronic or intergenic regions (894 out of 1,335 NFRs compared to 336 mRNA promoters). Thus, I-BRD9 treatment leads to a preferential downregulation of enhancer transcription throughout the HeLa EGF induction time course. Notably, 86% of downregulated FANTOM5 enhancers were depleted of GFP-GLTSCR1L by I-BRD9. Reciprocally, GFP-GLTSCR1L depleted intergenic/intronic NFRs in general (Wilcoxon signed-rank test, P = 8.9×10^−5^ and P = 2.1×10^−4^ for intergenic and intronic NFRs, respectively, Fig. 4d) and FANTOM5 enhancers in particular (Wilcoxon signed-rank test, P = 3.7×10^−9^) were associated with downregulated expression. These results indicate that transcription of enhancers, but not mRNA genes, is affected by H3K27ac-recognition by GBAF through the bromodomain of BRD9.

Although mRNA expression levels were, to a large extent, not affected by BRD9 inhibition, we hypothesized that downregulation of enhancer expression levels reflects reduced enhancer regulatory activities. To test this hypothesis, we compared expression levels of putative enhancer-promoter pairs contained within the same topologically associating domains (TADs) of HeLa cells (Rao et al. 2014). Indeed, promoters within TADs containing at least one differentially downregulated enhancer showed significantly larger downregulation upon I-BRD9 treatment in all time points when compared to promoters in TADs not containing any downregulated enhancer (Fig. 4e). These results indicate that GBAF is important not only for enhancer transcription but also for enhancer regulatory activities.

We posited that GBAF may have a role in forming or maintaining permissive chromatin at enhancers. At putative intronic/intergenic enhancers associated with transcriptional downregulation and GFP-GLTSCR1L depletion by I-BRD9, we noted a significant reduction in chromatin accessibility (Wilcoxon signed-rank paired test, P = 5.9×10^−9^, Supplementary Fig. 5e). Taken together, our data indicate that I-BRD9 leads to a reduced rate of eRNA transcription at GLTSCR1L-depleted enhancers and a downregulation of their target genes. We hypothesize that this is the consequence of reduced GBAF chromatin remodeling activity at enhancers caused by disrupted recognition of H3K27ac by BRD9 through inhibition of its bromodomain.

## Discussion

Although H3K27ac is frequently associated with active enhancers, its role in enhancer activity and its protein interaction environment have remained elusive. In this study, we applied a SILAC histone-peptide pulldown MS approach to identify protein complexes that interact with H3K27ac alone and in combination with H3K23ac. This approach allowed us to assess acetyl-lysine interaction preferences and to identify the SEC and GBAF complexes as major interactors of H3K27ac. Specifically, we identified H3K27ac interactions with both BRD9 and GLTSCR1L, definitive subunits of the GBAF complex (Alpsoy & Dykhuizen 2018), indicating a H3K27ac-associated function of GBAF.

Our systematic investigation of the subunits of GBAF revealed insights into their putative functions. Our data refines the composition of GBAF (Alpsoy & Dykhuizen 2018) by adding BCL7A/B/C as an additional subunit. Interestingly, none of the subunits of the GBAF complex contains known DNA binding domains, unlike those of canonical BAF/PBAF complexes (e.g. ARID2/ARID1/1a, PBRM1, SMARCE1, SMARCB1, PHF10 (Supplementary Table 1)). Hence, the GBAF complex has only two functional chromatin interaction domains, the bromodomains of SMARCA2/4 and BRD9. Due to this specific feature, we hypothesize that targeting of GBAF to chromatin is more sensitive to the recognition of acetylated histones (H3K27ac), than the canonical SWI/SNF complexes that contain sequence-independent DNA-binding subunits. Investigation of the function of the GiBAF-domain of GLTSCR1L revealed that it is capable of binding all inferred GLTSCR1L- and BRD9-interacting proteins as well as BRD4 and AFF1. We also found that the GiBAF-domain is responsible for the nuclear localization of GLTSCR1L. Other parts of the GLTSCR1L protein may be involved in Pol II interactions or pre-initiation complex formation, as indicated by the enrichment of TFIID components in GFP pulldown MS results of a GiBAF-domain deletion mutant.

To investigate the role of acetyl-lysine recognition by BRD9, we made use of I-BRD9, a selective inhibitor of the BRD9 bromodomain. I-BRD9 is a useful tool since it doesn’t impair any function of BRD9 beyond its acetyl-lysine recognition (Theodoulou et al. 2016). Our histone-peptide pulldown immunoblot experiments with I-BRD9 revealed that GLTSCR1L chromatin interaction is mediated by the bromodomain of BRD9 and is H3K27/23ac dependent. Upon I-BRD9 treatment, BRD9 lost specificity in acetyl-lysine recognition but remained bound to histone peptides. This is likely due to its participation in several BAF subcomplexes, where binding is mediated by acetyl-lysine recognition modules of other BAF subunits, such as the bromodomain of SMARCA2/4. Concomitantly with these proteomic interaction results, GLTSCR1L and BRD9 colocalized at NFRs flanked by H3K27ac. We observed an enrichment of BRD9 and GLTSCR1L binding at intronic and intergenic NFRs and to a lesser extent at gene promoters, and used I-BRD9 to verify the role of the BRD9 bromodomain in the targeting of GBAF complex to chromatin. Upon I-BRD9 treatment, in agreement with our histone peptide pulldown immunoblot results, GFP-GLTSCR1L was observed to dislocate from H3K27ac-associated intronic/intergenic chromatin and, in particular, from enhancers.

The observed loss of GLTSCR1L from enhancers upon BRD9 inhibition indicates that GBAF has a specific role at enhancers. This hypothesis is supported by a decrease in chromatin accessibility at GLTSCR1L-depleted enhancers and an overall downregulation of enhancer transcription initiation events in I-BRD9-treated HeLa cells stimulated with EGF. While overall mRNA expression levels were not affected by I-BRD9, genes residing within the same TADs as downregulated enhancers tended to follow the same trend. Therefore, we propose that GBAF is important for both enhancer transcription and enhancer regulatory activity. However, no downregulation was observed for *NR4A1* mRNA expression levels upon downregulation of eRNAs from its cognate enhancers by I-BRD9. This may be a consequence of enhancer-independent EGF stimulation of *NR4A1* gene expression by promoter-proximal events. Nevertheless, as the timing of events in the communication between enhancers and promoters are poorly understood, we do not rule out that the *NR4A1* locus is in an alternative mode of regulation.

While several of our experiments point at an enhancer-associated function of GBAF, further investigations are needed to assess its exact role and the associated mechanisms. Although we observed a minor but significant decrease in chromatin accessibility at GLTSCR1L-depleted downregulated enhancers, we cannot rule out that this is a consequence of reduced Pol II activity. Precise nucleosome positioning methods are needed to verify abnormalities in the maintenance of the enhancer NFR and nucleosomal occupancy at sites of transcriptional enhancers in the absence of a functional GBAF complex. Numerous studies have observed enhancer-specific effects of different SWI/SNF complexes, which are mostly related to chromatin accessibility (Yu et al. 2013; Hodges et al. 2018; Nakayama et al. 2017; Alver et al. 2017; Wang et al. 2016). Given that GBAF is recruited to H3K27ac-marked enhancers, which are frequently associated with open chromatin, it is therefore possible that GBAF activity is secondary to chromatin opening and maintenance by other remodeling complexes. In line with its association with eRNA transcription, it is conceivable that GBAF has a role in the positioning of enhancer TSS-proximal nucleosomes. The interactions of GBAF with transcriptional regulators (such as SEC, BRD4, and TFIID) indicate that GBAF may have additional roles at enhancer TSSs in Pol II pre-initiation complex formation or in the recruitment or assembly of factors needed for pause release and transcriptional elongation. Due to the critical importance of correct enhancer activities in development and tissue homeostasis, we anticipate that dysfunction of GBAF or its targeting to chromatin may be connected with developmental abnormalities or complex diseases such as cancer.

## Methods

### Cell culture and inducible cell-line generation

For histone peptide pulldowns, HeLa S3 cells (ATCC) and IB10 mESCs (ATCC) were grown in DMEM SILAC medium without Lys and Arg (Silantes, 280001300), with dialyzed FBS (Silantes, 281000900) and supplemented either with light (R0K0) or heavy (R10K8) isotopes of lysine and arginine (Silantes, 282986440 Lys-0:HCl, 211604102 Lys-8:HCl, 282986444 Arg-0:HCl, 201604102 Arg-10:HCl). Additionally, IB10 mouse embryonic stem cells were cultured in the presence of 2i compounds as described (Kloet et al. 2016). For GFP pulldowns, ChIP-seq, and CAGE experiments, HeLa S3 cells were cultured in DMEM with 10% FBS, supplemented with glutamine and Pen-Strep. For induction and inhibition experiments, hEGF (Sigma-Aldrich, E9644; PeproTech, AF-100-15) was added to the culture medium at a concentration of 100 ng/ml and I-BRD9 (Tocris Bioscience, 5591) was added at a concentration of 10 μM.

To generate inducible cell lines, 3^*^10^5 HeLa S3 cells containing an integrated FRT site (van Nuland et al. 2013) were seeded on 6-well plates and transfected after 24 h with two vectors: 1) pOG44 expressing FLP recombinase under the control of human cytomegalovirus (CMV) promoter and carrying blasticidin selection marker; 2) pcDNA5/FRT containing a hygromycin selection marker, FRT recognition sites, and an N-terminal GFP fusion protein in frame with the gene of interest (GLTSCR1L, BRD9, BAF60a, BAF155, GLTSCR1L GiBAF-domain, or GLTSCR1L with GiBAF-deleted under the control of a doxycycline-inducible (TET-ON) CMV promoter. 16 h after transfection, selection media supplemented with 3 μg/ml blasticidin (Sigma) and 100 μg/ml hygromycin (Invitrogen) was applied to cells. Single colonies that remained after 10 days of selection were picked and propagated in single 30 mm plates and subsequently tested for the expression of desired proteins after induction with doxycycline (1 μg/ml).

### Nuclear extract and whole cell lysate preparation

Nuclear extracts were prepared according to (Dignam et al., 1983). Cells were trypsinized, harvested, washed twice with PBS, and centrifuged at 400 g for 5 min at 4 °C. Resuspended cell pellets were incubated for 10 min at 4 °C in five volumes of buffer A (10 mM HEPES-KOH, pH 7.9, 1.5 mM MgCl_2_, and 10 mM KCl), then pelleted at 400 g for 5 min at 4 °C. Cells were resuspended in two volumes of buffer A supplemented with protease inhibitors and 0.15% NP-40. Cells were homogenized by 30–40 strokes with a type B pestle in Dounce homogenizer. After homogenization, lysates were spun at 3,200 g for 15 min at 4 °C. Nuclear pellet was washed once with PBS and spun at 3,200 g for 5 min at 4 °C. Pellet was resuspended in two volumes of buffer C (420 mM NaCl, 20 mM HEPES-KOH, pH 7.9, 20% (v/v) glycerol, 2 mM MgCl_2_, and 0.2 mM EDTA) with 0.1% NP-40, protease inhibitors, and 0.5 mM dithiothreitol and incubated with rotation for 1 h at 4 °C, then spun at 20 000 g for 30 min at 4 °C. The supernatant (nuclear extract) was aliquoted and stored at −80 °C until further use.

Whole cell extracts were prepared by adding 5 cell pellet volumes of lysis buffer (0.5% NP40, 150 mM NaCl, 50 mM Tris pH 8.0, 10% Glycerol and 1 × Complete Protease Inhibitors). Cells were vortexed for 30 s and then incubated for 2 hr on a rotation wheel. Samples were then centrifuged at 4,000 g in a swinging bucket rotor for 30 min, after which soluble whole extracts were aliquoted and stored at -80 °C until further use.

### Histone-peptide pulldowns

Histone peptide pulldowns were performed as described in (Vermeulen 2012), with minor modifications. Briefly, biotinylated (modified and non-modified) histone H3 peptides (aa 15-36) were purchased from BIOSYNTAN GmbH. 50 μg of histone peptide per pull-down was incubated with 75 μl of MyOne Streptavidin C1 Dynabeads (Thermo Fisher, 65002) for 20 min at RT in peptide binding buffer (150 mM NaCl, 50 mM Tris–HCl, pH 8.0, 0.1% (v/v) NP40). Beads were washed three times with 1 ml protein binding buffer [150 mM NaCl, 50 mM Tris-HCl pH 8.0, 1% NP40, 0.5 mM DTT, 10 mM ZnCl_2_ and complete protease inhibitors – EDTA free (Roche)]. 350-700 μg of nuclear extract (diluted to 0.6 mg/ml) was incubated with immobilized histone peptides in protein binding buffer for two hours at 4 °C on a rotation wheel. Beads were washed five times with 1 ml of protein binding buffer containing 400 mM NaCl and finally twice with 1 ml of protein binding buffer. Beads from both pull-downs (with non-modified and modified peptide) were pooled, and bound proteins were eluted and visualized on 4 %–12 % SDS-PAA gradient gels (Invitrogen) by colloidal blue staining (Invitrogen). Lanes corresponding to Forward (heavy - modified peptide pulldown; light - non-modified peptide pulldown) and Reverse (light - modified peptide pulldown, heavy - non-modified peptide pulldown) experiments were divided into 6-8 pieces, sliced into small (~ 1 mm) fragments and then subjected to in-gel trypsin digestion essentially as described in (Shevchenko et al. 2007). Antibodies used for detection of histone peptide binding proteins from HeLa-FRT-GFP-GLTSCR1L nuclear extracts after SDS-PAAG using immunoblotting (Fig 3a; cells grown in non-SILAC medium) are listed in Supplementary Table 2.

### Mass spectrometry and data analysis of histone peptide pulldowns

After trypsin digestion of gel slices, peptides were extracted, desalted using StageTips (Rappsilber et al., 2003), and separated using an EASY-nLC (Proxeon) connected online to a LTQ-Orbitrap Fusion Tribrid mass spectrometer (Thermo Fisher Scientific). Scans were collected in data-dependent top speed mode with dynamic exclusion set at 60 seconds. Raw data were analyzed using MaxQuant version 1.5.1.0 with default settings and searched against the Uniprot mouse and human proteomes, release 2015_12. Analysis was performed using Perseus 1.5.5.3. After filtering, the mean value was calculated on the ratios from both forward and reverse experiments. Missing values were imputed with a normal distribution (settings: downshift = 1, window = 0) and scatter plots were made using R. Hierarchical clustering was used to generate heatmaps based on Euclidean distance (settings: linkage = average, number of clusters = 300, processing = k-means, iterations = 10, restarts = 1).

### GFP affinity purification mass spectrometry (AP MS-MS)

Nuclear extracts (NE) or whole cell extracts (WCE) from doxycycline-induced (for 16 hours) and non-induced cells were subjected to GFP-affinity enrichment using GFP nanotrap beads (Chromotek) in triplicate. For each pull-down, 1 mg of NE or 3 mg of WCE was incubated with 7.5 μl beads in incubation buffer (300 mM NaCl, 0.1 % NP-40, 0.5 mM DDT, 20 mM HEPES–KOH pH 7.9, 50 μg/ml ethidium bromide) in a total volume of 400 μl. Beads were washed twice with incubation buffer containing 0.5 % NP-40, twice with 1X PBS containing 0.5 % NP-40 and finally twice with 1X PBS. Affinity purified proteins were subject to on-bead trypsin digestion as described previously (Baymaz et al. 2014). Tryptic peptides were acidified and desalted using StageTips (Rappsilber et al. 2007) and separated with an online Easy-nLC 1000 (Thermo Scientific). Mass spectra were recorded on an LTQ-Orbitrap QExactive mass spectrometer (Thermo Fisher Scientific), selecting the top 10 most intense precursor ions for fragmentation, or on an LTQ-Orbitrap Fusion Tribrid mass spectrometer (Thermo Fisher Scientific). Scans were collected in data-dependent top speed mode with dynamic exclusion set at 60 seconds.

### LFQ peptide analysis and identification

Thermo RAW files from LFQ AP MS-MS were analyzed with MaxQuant version 1.5.1.0 using default settings and searching against the UniProt human proteome, release 2015_12. Additional options for match between runs, LFQ, and iBAQ were selected. The msVolcano Shiny application was used to produce volcano plots for GFP-affinity purification experiments (Singh et al. 2016). Stoichiometry calculations were produced essentially as described (Smits et al. 2013) using Perseus version 1.4.0.8 and in-house R scripts.

### Chromatin preparation

Attached HeLa cells were double cross-linked, first with DSG (ThermoFisher) for 40 min, then washed with PBS followed by treatment with 1% formaldehyde in PBS for 10 min at room temperature with gentle shaking. Crosslinking was quenched with the addition of 1/10 volume 1.25 M glycine. Cells were washed with PBS, then harvested by scraping in buffer B (20 mM HEPES, 0.25 % Triton X-100, 10 mM EDTA, and 0.5 mM EGTA). Cells were pelleted by centrifugation at 600 g for 5 min at 4 °C. Cell pellets were resuspended in buffer C (150 mM NaCl, 50 mM HEPES, 1 mM EDTA, and 0.5 mM EGTA) and rotated for 10 min at 4 °C. Cells were pelleted by centrifugation at 600 g for 5 min at 4 °C. The cell pellet was then resuspended in 1X incubation buffer (0.15% SDS, 1% Triton X-100, 150 mM NaCl, 1 mM EDTA, 0.5 mM EGTA, and 20 mM HEPES) at 15 million cells/mL. Cells were sheared in a Bioruptor Pico sonicator (Diagenode) at 4 °C with 7 cycles of 30 s on, 30 s off. Sonicated material was spun at 18,000 g for 10 min at 4 °C, then divided into aliquots and stored at −80 °C.

### Chromatin immunoprecipitation

10 million cells were used as an input material. Chromatin was incubated overnight at 4 °C in 1X incubation buffer (0.15 % SDS, 1 % Triton X-100, 150 mM NaCl, 1 mM EDTA, 0.5 mM EGTA, and 20 mM HEPES) supplemented with protease inhibitors and 0.1 % BSA. Antibody amounts and catalog numbers are listed in Supplementary Table 2. A 50:50 mix of Protein A and G Dynabeads (Invitrogen) were added the next day and incubated for 90 min. The beads were washed twice with wash buffer 1 (0.1 % SDS, 0.1 % sodium deoxycholate, 1 % Triton X-100, 150 mM NaCl, 1 mM EDTA, 0.5 mM EGTA, and 20 mM HEPES), once with wash buffer 2 (wash buffer 1 with 500 mM NaCl), once with wash buffer 3 (250 mM LiCl, 0.5% sodium deoxycholate, 0.5 % NP-40, 1 mM EDTA, 0.5 mM EGTA, and 20 mM HEPES), and twice with wash buffer 4 (1 mM EDTA, 0.5 mM EGTA, and 20 mM HEPES). After washing steps, beads were rotated for 20 min at room temperature in elution buffer (1 % SDS and 0.1 M NaHCO3). The supernatant was de-crosslinked with 200 mM NaCl and 100 μg/mL proteinase K for 4 h at 65 °C. De-crosslinked DNA was purified with MinElute PCR Purification columns (Qiagen). DNA amounts were determined with Qubit fluorometric quantification (ThermoFisher Scientific).

### Chromatin immunoprecipitation sequencing and data analysis

Libraries were prepared with a Kapa Hyper Prep Kit for Illumina sequencing (Kapa Biosystems) according to the manufacturer’s protocol with the following modifications. 5 ng DNA was used as input, with NEXTflex adapters (Bioo Scientific) and ten cycles of PCR amplification. Post-amplification cleanup was performed with QIAquick MinElute columns (Qiagen), and size selection was performed with an E-gel (300-bp fragments) (ThermoFisher Scientific). Size-selected samples were analyzed for purity with a High Sensitivity DNA Chip on a Bioanalyzer 2100 system (Agilent). Samples were sequenced on an Illumina HiSeq2000 or NextSeq500. Reads were mapped to the reference human genome hg19 with the Burrows–Wheeler Alignment tool (BWA), allowing one mismatch. Only uniquely mapped reads were used for data analysis and visualization. Each ChIP-seq bam file was first converted to tag align files using gawk and bedtools and peak calling was performed using MACS (version 2.1.0). Afterwards, to get a set of confident peaks, only peaks with Irreproducible Discovery Rate (IDR) < 0.05 were kept.

### CAGE library preparation, sequencing and mapping

HeLa S3 cells were grown on 60 mm plates and treated with DMSO or I-BRD9 (10 μM) 6 h prior to induction with EGF (100 ng/ml) for 30 min. Total RNA was isolated using TRI Reagent^®^ (Ambion) according to manufacturer’s recommendations. RNA from each of the biological triplicates were quality controlled using a Bioanalyzer. RIN scores were between 9.6 and 10. CAGE libraries were prepared using the protocol by (Takahashi et al. 2012) with an input of 3 μg of total RNA. Prior to sequencing, four CAGE libraries with different barcodes were pooled and applied to the same sequencing lane. Libraries were sequenced on a Illumina HiSeq 2000 at the National High-Throughput DNA Sequencing Centre, University of Copenhagen. To compensate for the low complexity of 5′ ends in the CAGE libraries, 30% Phi-X spike-ins were added to each sequencing lane, as recommended by Illumina. CAGE reads were assigned to their respective originating sample according to identically matching barcodes. Using the FASTX Toolkit, assigned reads were 5′-end trimmed to remove linker sequences (9+2 bp to account for the CAGE protocol G-bias), 3′-end trimmed to a length of 25 bp, and filtered for a minimum sequencing quality of Q30 in 50% of the bases. Reads matching to reference rRNA sequences were discarded using rRNAdust. Mapping to the human genome (hg19) was performed using BWA (version 0.7.10). Only the 5’ ends of mapped reads were considered in subsequent analyses.

### ATAC-seq library preparation and processing

ATAC-seq was performed on approximately 50,000 cells as described in (Buenrostro et al. 2015) with three modifications. First, the total volume of the tagmentation reaction with in-house made Tn5 enzyme was halved. Second, the tagmentation reaction was stopped with 44 mM EDTA, 131 mM NaCl, 0.3% SDS, and 600 μg/ml proteinase K. Lastly, a reverse-phase 0.65× SPRI beads (Ampure) DNA purification was done after the first PCR. Libraries were sequenced on an Illumina HiSeq 2000. Paired-end 50-bp sequencing reads were aligned to hg19 with BWA (version 0.7.10) allowing one mismatch. To produce a consensus set of ATAC-seq peaks, bam files were first converted to tag align files using gawk and bedtools and peak calling was performed on all the samples (pooled and individually) using MACS (version 2.1.0) after shifting tag alignments (+4 bp for plus strand and -5bp for minus strand) to account for TN5 insertion. Afterwards, narrow peaks were identified from the pooled list overlapping at least two individual peak lists by at least 50% of bps (FDR < 1%). This resulted in a preliminary list of 229,091 peaks.

### Open chromatin loci as focus points for transcription initiation and expression quantification

Open chromatin loci (also referred to as NFRs) were used as focus points for characterizing transcription initiation events as described elsewhere (Andersson et al. 2014), with minor modifications. Instead of focusing on DNase-seq signal summits, center points were defined from ATAC-seq peak signal summits. Open chromatin loci were filtered to not overlap any other open chromatin loci strand-specifically with respect to these windows. This resulted in a final set of 213,017 well-defined open chromatin loci. NFR-associated expression were quantified by counting of CAGE tags in genomic windows of 300 bp immediately flanking ATAC-seq peak summits. An average of 79% of all CAGE tags were covered by the filtered set of open chromatin loci.

For robust assessment of lowly expressed loci, CAGE genomic background noise levels were estimated as described elsewhere (Rennie et al. 2018). First, the CAGE mappability of the hg19 reference genome was calculated by mapping each 25-sized subsequence of the reference genome back to itself, using the same mapping approach as for real CAGE data. Then, the number of CAGE 5’ ends from each CAGE library mapping to each of two strand-specific genomic windows genomic regions of size 300 bp was quantified, similarly to the expression quantification of NFRs (above). Genomic windows were required to be uniquely mappable (as determined by the mappability track) in at least 50% of its potential TSS positions (unique bps). Regions that were proximal (within 500bp) of GENCODE (v19) gene TSSs, transcript ends, or midpoints of ENCODE DHSs (ENCODE January 2011 integration data), or overlapping GENCODE gene exons were discarded. Based on the empirical distribution of CAGE expression noise from annotation-distal genomic regions, the 99th percentile for each library was used as a threshold to call regions significantly expressed in subsequent analyses, if fulfilled in at least 2 out of 3 replicates. This resulted in a set of 20,303 and 22,771 open chromatin loci expressed in the I-BRD9 and DMSO time courses respectively.

Differential expression analysis was performed on all expressed open chromatin loci using DESeq2. To find differential dynamic open chromatin, comparisons were made between all time points T>0 and T=0. This resulted in a total of 1,709 differentially dynamic open chromatin loci (FDR<5%, |log_2_ FC| > 1) in I-BRD9 and 1,922 in DMSO (control) time course. Differential expression analysis was also performed between conditions on the same time points, resulting in 5,633 open chromatin loci with expression differences in at least one time point, of which 710 were also differentially dynamic.

## Data availability

Sequencing data (ChIP-seq, ATAC-seq, CAGE) generated in this study have been deposited in GEO under accession number GSE121351. Proteomics data generated in this study have been deposited in the PRIDE archive of the ProteomeXchange consortium under accession number PXD011376.

## Acknowledgements

We would like to thank Ragnhild Eskeland for rewarding discussions and for kindly sharing the pEF1neo-TY1-tag plasmid. We would also like to thank members of the Vermeulen, Aasland, and Andersson labs for insightful comments and suggestions. Research in Vermeulen lab was supported by 4DCellFate within the European Commission’s Seventh Framework Program (277899), and the European Research Council (StG no. 309384)., The Vermeulen lab is part of the Oncode Institute, which is partly funded by the Dutch Cancer Society (KWF). Research in the Aasland lab was supported by the Norwegian Cancer Society (#197477). Research in the Andersson lab was supported by Independent Research Fund Denmark (6108-00038B) and the European Research Council under the European Union’s Horizon 2020 research and innovation programme (StG no. 638173).

## Author contributions

K.J. and R.Aa conceived the project. M.V., R.Aa, and R.An. supervised the project. K.J., S.L.K., M.V., and R.Aa. designed proteomics experiments and K.J., M.V., R.Aa., and R.An. designed sequencing experiments. K.J. and S.L.K. performed mass spectrometry experiments. S.V. performed qPCR experiments and MS data validation by immunoblotting. S.A.S. performed immunofluorescence experiments. S.L.K. performed ATAC-seq experiments. S.V. and C.D.V. performed CAGE experiments. K.J. and S.L.K. performed ChIP-seq experiments. K.J. and S.L.K. analyzed mass spectrometry data. N.A. analyzed ATAC, ChIP, and CAGE sequencing data and performed integrative bioinformatics analyses. K.J. and R.An. wrote the manuscript with input from all authors.

## Footnotes

1 GLTSCR1L and GLTSCR are paralogs of a group of proteins found in vertebrates. They are also known by the names BICRAL and BICRA (UniProt IDs Q6AI39 and Q9NZM4, respectively). GLTSCR1L and GLTSCR contain a highly conserved domain, which we refer to as ‘GiBAF’ (<u>G</u>LTSCR1/1L domain interacting with <u>BAF</u> complex; Supplementary Fig. 2).

